# Purine nucleobases enhance CD8^+^ T cell effector function

**DOI:** 10.1101/2025.08.14.669899

**Authors:** Jakob-Wendelin Genger, Ben Haladik, Alexander Lercher, Benedikt Agerer, Carolina Mangana, Fabian Amman, Csilla Viczenczova, Theresa Hohl, Anna Hofmann, Maximilian Koblischke, Bernhard Kratzer, Wentao Li, Kerstin Zinober, Felix Kartnig, Christina Schüller, Anna Koren, Jung-Ming G. Lin, Barbara B. Maier, Christoph Bock, Robert Kralovics, Rafael J. Argüello, Judith H. Aberle, Thomas Reiberger, Winfried F. Pickl, Giulio Superti-Furga, Kristaps Klavins, J. Thomas Hannich, Stefan Kubicek, Andreas Bergthaler

**Author notes:** Contact info.

## Abstract

During antiviral immune responses, activated immune cells remodel metabolic pathways towards uptake and utilization of biosynthetic and bioenergetic metabolites. Concurrently, viral infections alter metabolic environments, impacting metabolite availability for the establishment of an effective immune response. Here, we integrated *in vivo* metabolomics data from murine and human viral infections with *in vitro* metabolite screens, identifying purine nucleobases as novel *immunometabolites* that enhance CD8^+^ T cell effector function. We found that CD8^+^ T cells can switch from resource-intensive purine *de novo* synthesis to purine salvage pathway, to produce nucleotides from purine nucleobases. This strategy of metabolic adaptation allows diversion of biosynthetic and bioenergetic resources towards enhancing effector molecule production. Our findings unveil an adaptation strategy by CD8^+^ T cells to manage metabolic challenges in dynamic organismal environments and suggest pharmacological targets in purine metabolism as potential targets for immunotherapy.

**Graphical Abstract:** 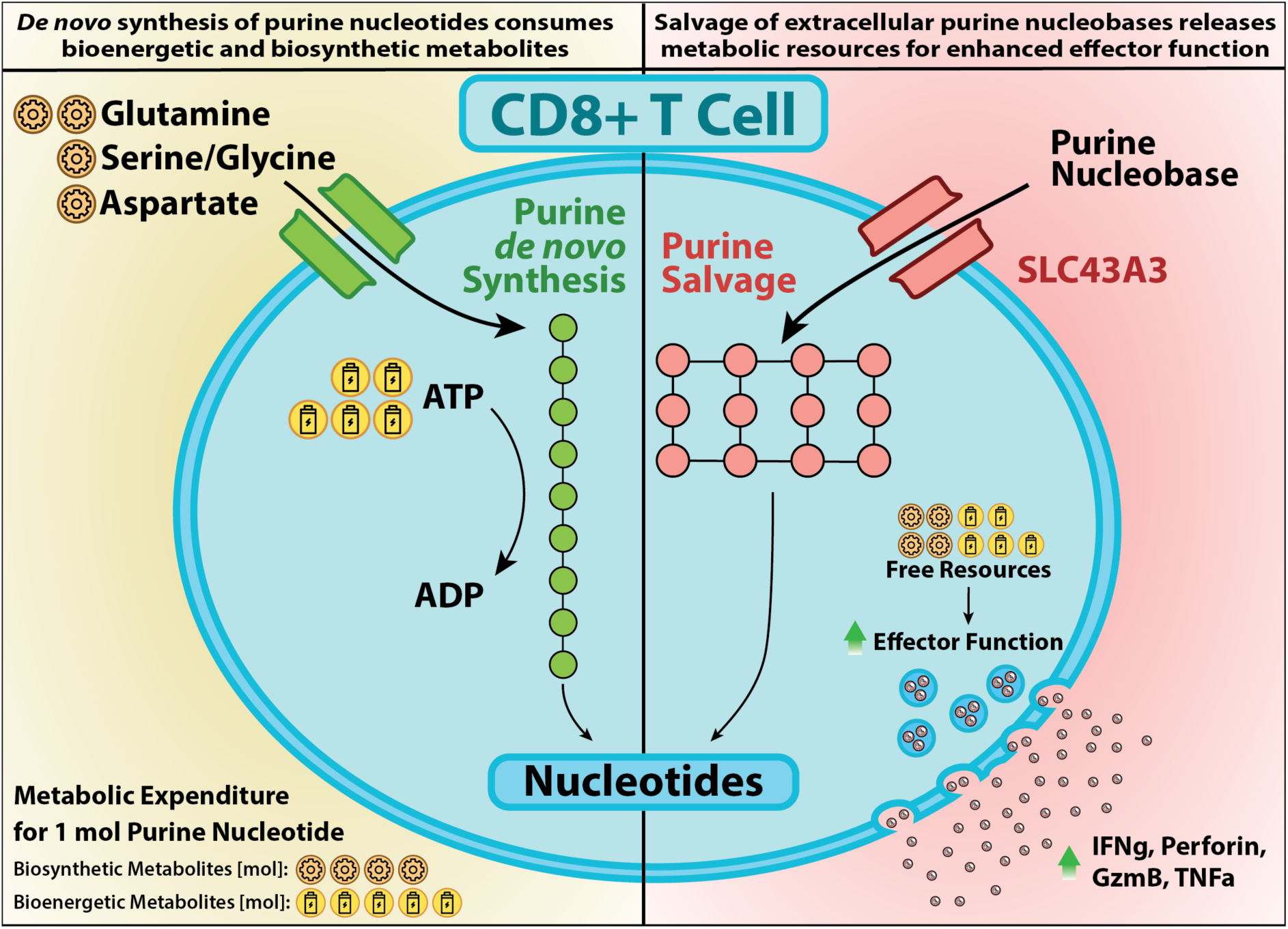

Instead of producing nucleotides via purine *de novo* synthesis, CD8^+^ T cells can import and utilize purine nucleobases via the purine salvage pathway to divert bioenergetic and biosynthetic resources towards effector function. By shifting from purine *de novo* synthesis to the purine salvage pathway, cells save significant resources: 5 moles of the key bioenergetic metabolite ATP, and biosynthetic metabolites including 2 moles of glutamine, 1 mole each of serine or glycine, and 1 mole of aspartate.

## Introduction

Upon activation, immune cells typically undergo enhanced proliferation, migration, and secretion of effector molecules. However, clonal expansion and effector molecule production are metabolically demanding^1–6^. Metabolites play a pivotal role in modulating immunity through various mechanisms: Firstly, as bioenergetic fuels that meet cellular energy requirements; secondly, as biosynthetic substrates essential for generating effector molecules and maintaining cell physiology, including organelle remodeling; and thirdly, as molecules facilitating both intra-and intercellular signaling^3,5,7^. These metabolites build a bridge between metabolism and immunity and were termed *immunometabolites*^8^.

An increasing number of publications elucidated the metabolic requirements for the establishment of effective T cell responses. Metabolites, including glucose and several amino acids, are indispensable as bioenergetic and biosynthetic substrates for T cell activation and effector function^3,5,9–13^. Beyond their role as metabolic fuel, specific metabolites such as kynurenine and short-chain fatty acids exert regulatory effects on T cells as signaling molecules or substrates for epigenetic modification reactions ^3,5,14,15^. Therefore, the availability of these immunometabolites modulates T cell responses^3,5,16^.

Changes in local and systemic metabolic environments have been described in various settings of inflammation, such as the tumor microenvironment and infection^3^. These metabolic changes result from a variety of processes induced by immune mechanisms or the invading pathogen^3,17^. Viruses, characterized by diverse tissue tropisms, virulence, and varying infection kinetics, are metabolic engineers that reprogram their host’s metabolism to facilitate their replication^17^. CD8^+^ T cells are a central component of the defense against intracellular pathogens, such as bacteria and viruses, and tumor cells^18^. However, how infection-associated changes in metabolic environments affect the establishment of CD8^+^ T cell responses remains understudied.

Here, we explored the biological mechanisms of metabolic modulation of antiviral immunity and utilized different viral infections as tools to identify novel immunometabolites that shape CD8^+^ T cell responses. First, we generated an *in vivo* infection metabolomics dataset to systematically investigate the metabolic alterations associated with various viral infections. Then, we designed a metabolite screening approach to identify altered metabolites that modulate CD8^+^ T cell proliferation, activation, and effector function *in vitro*. Our results revealed purine nucleobases as immunometabolites that are recycled through the purine salvage pathway for the biosynthesis of nucleotides. This mechanism spares bioenergetic and biosynthetic resources of CD8^+^ T cell, resulting in enhanced effector function (see **Graphical Abstract**). Thus, this study_shines new light on the role of purine metabolism as central immunometabolic hub for the establishment of effective CD8^+^ T cell responses.

## Results

### Serum metabolomics analysis reveals dynamic patterns in serum metabolite composition during viral infections

To unravel how the systemic availability of metabolites changes during viral infections, we tailored a targeted metabolomics panel for the quantification of 201 metabolites. This targeted panel comprised key positions in major metabolic pathways ranging from glycolysis, TCA cycle and purine metabolism to biogenic amines and amino acids. We employed this analytical panel to assess serum samples from a range of experimental murine viral infection models comprising the orthomyxovirus Influenza A Virus strain Puerto Rico 8 (IAV PR8), the arenavirus lymphocytic choriomeningitis virus (LCMV) strains Clone 13 (Cl13) and Armstrong (ARM), and the coronavirus Murine Hepatitis Virus A59 (MHV). These models were selected to encompass both, acute and chronic infections, as well as viruses with systemic and localized pathology. The LCMV models, pivotal in viral immunology and T cell biology, offer essential tools and resources to study various aspects of antiviral T cell responses, making the two strains invaluable for investigating distinct facets of these immune responses^19–23^. LCMV ARM causes an acute infection which is cleared by a vigorous CD8^+^ T cell response within 8-10 days^23^. LCMV Cl13 establishes a chronic infection accompanied by immunopathological manifestations such as T cell-mediated hepatitis and subverts the immune response via induction of T cell exhaustion^24–27^. IAV PR8 causes an acute respiratory infection restricted to the lungs^28^. We also included the coronavirus MHV A59, a cytopathic, neuro-and hepatotropic coronavirus model that induces acute hepatitis^29^. Upon infection, we collected sera from mice at 2 and 8 days post infection (dpi), approximating the peak of the innate and adaptive immune responses, respectively.

We noted the significant differential abundance of 151 detectable serum metabolites across multiple major metabolic pathways at 8 dpi (**Fig. 1a, Extended Data Fig. 1a-b and Extended Data Table 1**). The most pronounced alterations were observed in TCA cycle metabolites and purine metabolism, specifically hypoxanthine, xanthine, inosine and adenosine (**Fig. 1a and Extended Data Fig. 1a**). LCMV Cl13 exhibited strong metabolic correlation with LCMV ARM at both 2 and 8 dpi with purine metabolites among the most downregulated metabolites during the peak of the adaptive immune response (**Fig. 1a-b and Extended Data Fig. 1c**). Interestingly, the two infection models LCMV Cl13 and MHV showed correlated metabolic alterations only during the peak of the innate immune response 2 dpi but diverged by 8 dpi (**Fig. 1c and Extended Data Fig. 1c**). Principal component analysis (PCA) further substantiated that the different courses of infection are distinguishable by their metabolic trajectories (**Fig. 1c**). Moreover, the results highlighted a substantial influence of the progression of the infections and their corresponding immune responses on host systemic metabolism (**Fig. 1c**). This implies that metabolic changes are stronger during the peak of the adaptive immune response than during the peak of the innate immune response across virus models, possibly also reflective of ongoing tissue pathology and organismal metabolic changes. Metabolite set enrichment analysis ranked purine metabolism and related metabolic pathways, such as the pentose phosphate pathway, among the top 10 most infection-associated enriched KEGG metabolite sets 8 dpi (**Fig. 1d and Extended Data Fig. 1d**). Together, these results suggest temporal and pathogen-specific nodes of systemic metabolic changes during infection.

**Figure 1:**
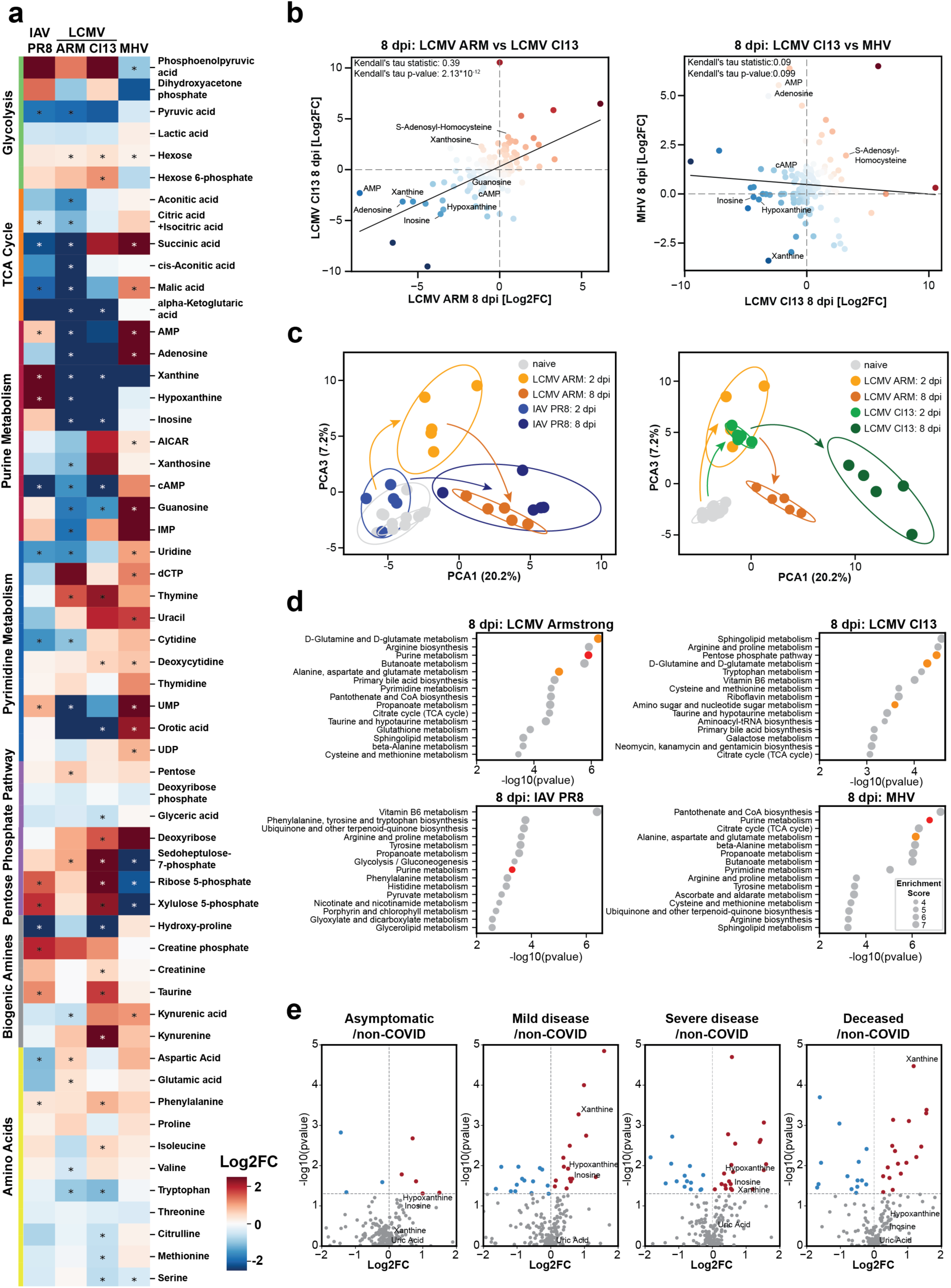
Comprehensive analysis of systemic metabolism reveals dynamic patterns in serum metabolite concentrations during viral infections. **a)** Longitudinal profiling of serum metabolite changes 8 dpi in five age-and sex-matched mice per group infected with LCMV Armstrong (LCMV ARM), LCMV Cl13, Influenza A virus Puerto Rico 8 (IAV PR8) or murine hepatitis virus (MHV). Statistical analysis: Longitudinal metabolomics measurements were analyzed with paired t-tests (naïve vs 8 dpi) with two-stage step-up Benjamini-Hochberg procedure to control the FDR (alpha = 0.1). **b)** Correlation matrix of serum metabolite changes 8 dpi with linear regression line. **c)** Principal component analysis (PCA) visualizing the divergence and convergence of metabolic trajectories among LCMV ARM, LCMV Cl13, and IAV PR8-infected mice. **d)** Metabolite set enrichment analysis indicating perturbed metabolic pathways during infection with LCMV ARM, LCMV Cl13, IAV PR8, and MHV at 8 dpi. Purine metabolism is emphasized in red, with associated metabolic pathways marked in yellow. **e)** Serum metabolomic profiling across different clinical severity stages of COVID-19, based on data adapted from Valdés *et al.*^30^

We aimed to validate our findings on infection-associated metabolic changes in different murine models by clinical metabolomics data obtained from patients with hepatotropic and respiratory viral infections. To this end, we analyzed serum samples from a defined patient cohort with hepatitis C virus (HCV) infection at timepoints pre-and post-curative treatment^30^. In line with our observations in the murine infection models, purine metabolism ranked among the most significantly enriched metabolite sets during HCV infection with xanthine and inosine among the most differentially regulated metabolites **(Extended Data Fig. 1e-f)**. Moreover, we examined available datasets from COVID-19 patients across different severity stages^30^. Interestingly, we observed that the serum of COVID-19 patients with varying disease severity, from mild to fatal outcomes, displayed comparable increases in purine metabolites such as xanthine and hypoxanthine, analogous to what we observed during IAV PR8 infection in mice (**Fig. 1e**)^30^.

We observed that metabolic changes were more pronounced during the peak of the adaptive than the innate immune response. Antiviral CD8^+^ T cells are a central part of the adaptive immune response against viruses. These rapidly proliferating effector CD8^+^ T cells have a high demand for bioenergetic and biosynthetic metabolites^5^. We, thus, wanted to investigate how the activity of metabolically demanding CD8^+^ T cells is affected by the drastically changing metabolic environments in an infected host. Upon reanalysis of available transcriptomics data from LCMV-specific CD8^+^ T cells, we consistently identified purine metabolism as top hit among the most prominent differentially regulated metabolic pathways during the establishment of adaptive antiviral CD8^+^ T cell response (**Extended Data Fig. 1g**)^31^.

In summary, our metabolomics serum analyses across different viral infections revealed a plethora of infection-associated metabolic changes that may be indicative of the individual viral pathogenic mechanisms, host responses, and/or stages of the immune responses. Concurrently, CD8^+^ T cells undergo vast metabolic remodeling during the establishment of an antiviral adaptive immune response.

### Purine metabolism is a key metabolic node of CD8^+^T cell effector function

Next, we set out to assess which of the infection-associated differentially regulated serum metabolites could play a role in modulating the establishment of antiviral CD8^+^ T cell responses. We established a high-throughput image-based screening approach to elucidate the effects of target immunometabolites on CD8^+^ effector function *in vitro* (**Fig. 2a**). Specifically, we interrogated the effect of 58 differentially regulated metabolites on proliferation, activation and effector function by quantifying cell counts, expression of CD44 and interferon-γ (IFNγ) respectively (**Fig. 2b, Extended Data Fig. 2a and Extended Data Table 2**). Using this workflow, we identified the purine nucleobases hypoxanthine and adenine as enhancers of CD8^+^ T cell effector function in a dose-dependent fashion, while proliferation remained unaffected (**Fig. 2c-d and Extended Data Fig. 2b-d**). We corroborated these results through quantification of IFNγ concentrations in cell supernatants by ELISA (**Extended Data Fig. 2e**). Further, our transcriptional analysis of CD8+ T cells treated with hypoxanthine or adenine confirmed the upregulation of several immune pathways related to cytokine production and inflammation by gene set enrichment analysis **(Fig. 2e)**.

**Figure 2:**
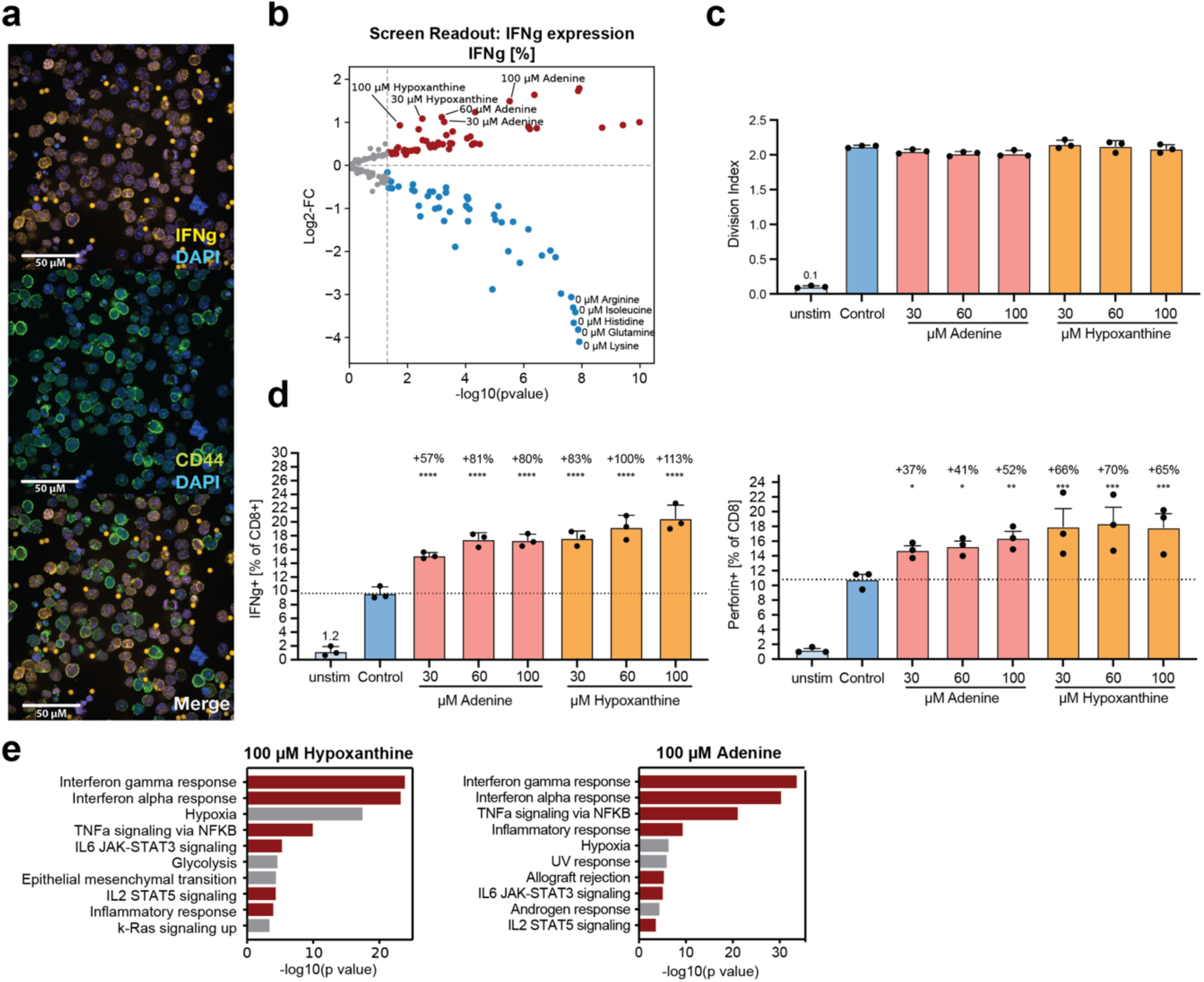
Purine metabolism is a key metabolic node for CD8^+^ T cell effector function. **a)** Representative immunofluorescence images for the metabolite screen setup. **b)** Results of Interferon-γ (IFNγ) expression readout in high-throughput metabolite screen setup. Unstimulated and staining controls were removed from the visualization for enhanced clarity. Statistical analysis: Independent t-test with Benjamini-Hochberg procedure to control the FDR (alpha = 0.1) with sample size n = 6. **c-d)** Flow cytometry-based validation of metabolite screen outcomes examining the impact of purine nucleobases on **c)** cellular proliferation and **d)** effector function. The graph is a representative example of three independent experiments. Data are represented as mean ± SD with sample size n = 3. **e)** Gene set enrichment analysis of genes differentially regulated with supplementation of 100 µM hypoxanthine or 100 µM adenine compared to untreated control. The sample size is n = 3.

As control, we also tested the corresponding nucleosides, inosine, and adenosine as well as the downstream products of purine nucleobases in purine metabolism, xanthine and uric acid in the same concentration range (**Extended Data Fig. 2f-g**). These experiments found no discernible effect on the effector function of CD8^+^ T cells, suggesting that observed effects on CD8^+^ T cells are specific to purine nucleobases.

### Salvaging purine nucleobases spares biosynthetic and bioenergetic resources in CD8^+^ T cells for enhanced effector function

To further elucidate the role of purine metabolism during CD8^+^ T cell activation, we quantified the intracellular concentrations of metabolites by targeted metabolomics. Among the intracellular metabolites that were elevated during T cell activation, 24 out of 30 purine metabolites in our assay panel exhibited significant upregulation 48 hours post-stimulation, and 9 remained significantly elevated 72 hours post-stimulation (**Fig. 3a and Extended Data Table 3**). Overall, purine metabolites were among the most prominently upregulated intracellular compounds, which supports the notion that the management of cellular purine metabolite pool is important to CD8^+^ T cell activation.

**Figure 3:**
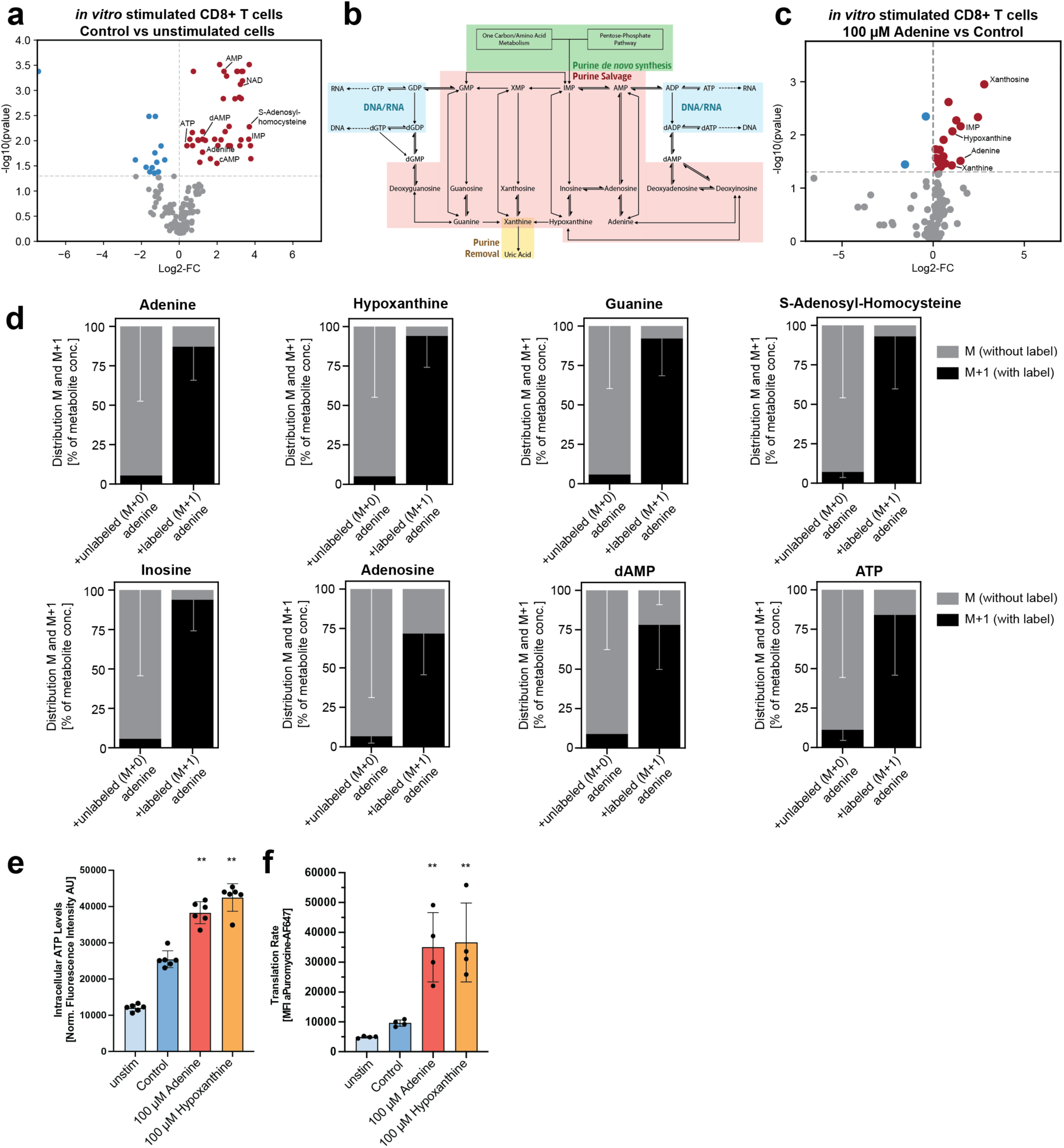
Salvaging purine nucleobases spares biosynthetic and bioenergetic resources in CD8^+^ T cells for enhanced effector function. **a)** Changes in intracellular metabolite concentrations in CD8^+^ T cells after 72h *in vitro* stimulation with plate-bound anti-CD3/anti-CD28 antibodies. Statistical analysis: Independent t-test with two-stage step-up Benjamini-Hochberg procedure to control the FDR (alpha = 0.1) with sample size n = 3. Data are represented as percentage of total with standard deviation. **b)** Schematic overview of the metabolic pathways in purine metabolism. **c)** Intracellular metabolite concentrations in CD8^+^ T Cells treated with 100 µM adenine versus untreated controls 72 hours post-stimulation. Statistical analysis: Independent t-test. **d)** Mass spectrometry-based analysis of the distribution of heavy isotope-labelled C atoms in cellular purine metabolites. CD8^+^ T cells were stimulated *in vitro* in cell culture media with dialyzed FBS for 72 hours with supplements of 8-^13^C heavy-isotope labelled adenine before measurement. **e)** Measurement of intracellular ATP levels using the CellTiter-Glo^®^ method in *in vitro* stimulated CD8+ T cells normalized by cell viability. Statistical analysis: Independent t-test. **f)** Translation rate in *in vitro* stimulated CD8+ T cells determined via puromycin incorporation using the SCENITH method. Statistical analysis: Independent t-test. **a,c,d)** The sample size is n=3. **e-f)** Data are represented as means ± SD with sample size **e)** n = 3 or **f)** n = 4, respectively.

Cellular purine metabolism consists of two consecutive pathways: purine *de novo* synthesis and purine salvage (**Fig. 3b**)^32,33^. The resource-intensive purine *de novo* synthesis pathway synthesizes new purines from glutamine, serine or glycine, and aspartate, utilizing 5 ATP molecules to produce each molecule of the end product, inosine monophosphate (IMP). It has been demonstrated to play a critical role for satisfying the metabolic demand for nucleotides to drive proliferation during T cell activation^9^. The purine salvage pathway maintains and balances intracellular purine pools of nucleobases, nucleosides, and nucleotides by facilitating conversions among hypoxanthine-, guanine-, and adenine-based purines such as IMP. This resource-efficient pathway enables the synthesis of nucleotides from nucleobases recycled from DNA and RNA, or from extracellular purine sources, independently of purine *de novo* synthesis. This process can conserve cellular biosynthetic and bioenergetic resources. We hypothesized that CD8^+^ T cells are metabolically flexible to feed purine nucleobases into the purine pool via the purine salvage pathway, thereby remodeling their purine metabolism towards optimized resource efficiency and enhancing effector function^33^.

We tested the ability of CD8+ T cells to actively import and utilize extracellular purine nucleobases via the purine salvage pathway by supplementing with 100 µM adenine during activation. This supplementation resulted in elevated concentrations of related purines and IMP, suggesting the capability of CD8^+^ T cells to effectively utilize these resources (**Fig. 3c**). To provide further evidence for the metabolic flexibility of CD8^+^ T cells to import and utilize extracellular purine nucleobases via the purine salvage pathway, we employed a metabolite tracing experiment using 100 µM of heavy-isotope-labeled 8-^13^C-adenine. As anticipated, the labeled ^13^C atom from adenine was incorporated in all measured purine metabolites. This includes nucleobases, nucleosides, nucleotides, and specialized purine metabolites such as the universal bioenergetic metabolite ATP, and S-adenosyl-homocysteine, the precursor of S-adenosyl-methionine, a donor of transmethylation reactions (**Fig. 3d**). As expected, the labelled ^13^C atom could not be found in metabolites upstream of or unrelated to purine metabolism such as cytidine, AICAR and serine (**Extended Data Fig. 3a**). Moreover, supplementation of adenine or hypoxanthine effectively increased ATP concentrations within CD8^+^ T cells, suggesting increased availability of bioenergetic resources (**Fig. 3e**). Using the Single Cell Energetic Metabolism by Profiling Translation Inhibition (SCENITH™) method, we also found that purine nucleobase supplementation increased the translation rate of CD8^+^ T cells (**Fig. 3f**)^34^. Taken together, these results demonstrate that the availability of purine nucleobases creates a metabolic environment where CD8+ T cells can switch from resource-intensive purine *de novo* synthesis to resource-efficient purine salvage which spares bioenergetic and biosynthetic resources to support translation and likely other cellular processes for enhanced effector function.

### Pharmacological perturbation of purine salvage pathway enhances CD8^+^ T cell effector function

To explore strategies to leverage intracellular purine metabolism for the modulation of CD8^+^ T cell effector function, we employed 6-mercaptopurine (6-MP), a purine analog that suppresses T cell proliferation via inhibition of synthesis and conversion of purine metabolites via the purine salvage pathway^33^. When we administered low doses of 6-MP (50 nM), which only modestly reduced the proliferative capacity of CD8+ T cells, we observed a significant increase in the production of the effector molecules IFNγ and perforin (**Fig. 4a,b and Extended Data Fig. 4a**). Quantification of IFNγ and TNFα concentrations in cell supernatants by ELISA further validated these observations (**Extended Data Fig. 4b**). Intracellular metabolite enrichment and transcriptional signatures were comparable to the supplementation of CD8^+^ T cells with 100 µM purine nucleobases (**Fig. 4c-d**). To further investigate whether analogs of 6-MP exhibit similar effects, we probed the purine metabolic network with additional compounds such as thioguanine and azathioprine. We found that these compounds showed the same effect on effector function to an extent comparable to that of 6-MP (**Fig. 4e-f and Extended Data Fig. 4c-d**).

**Figure 4:**
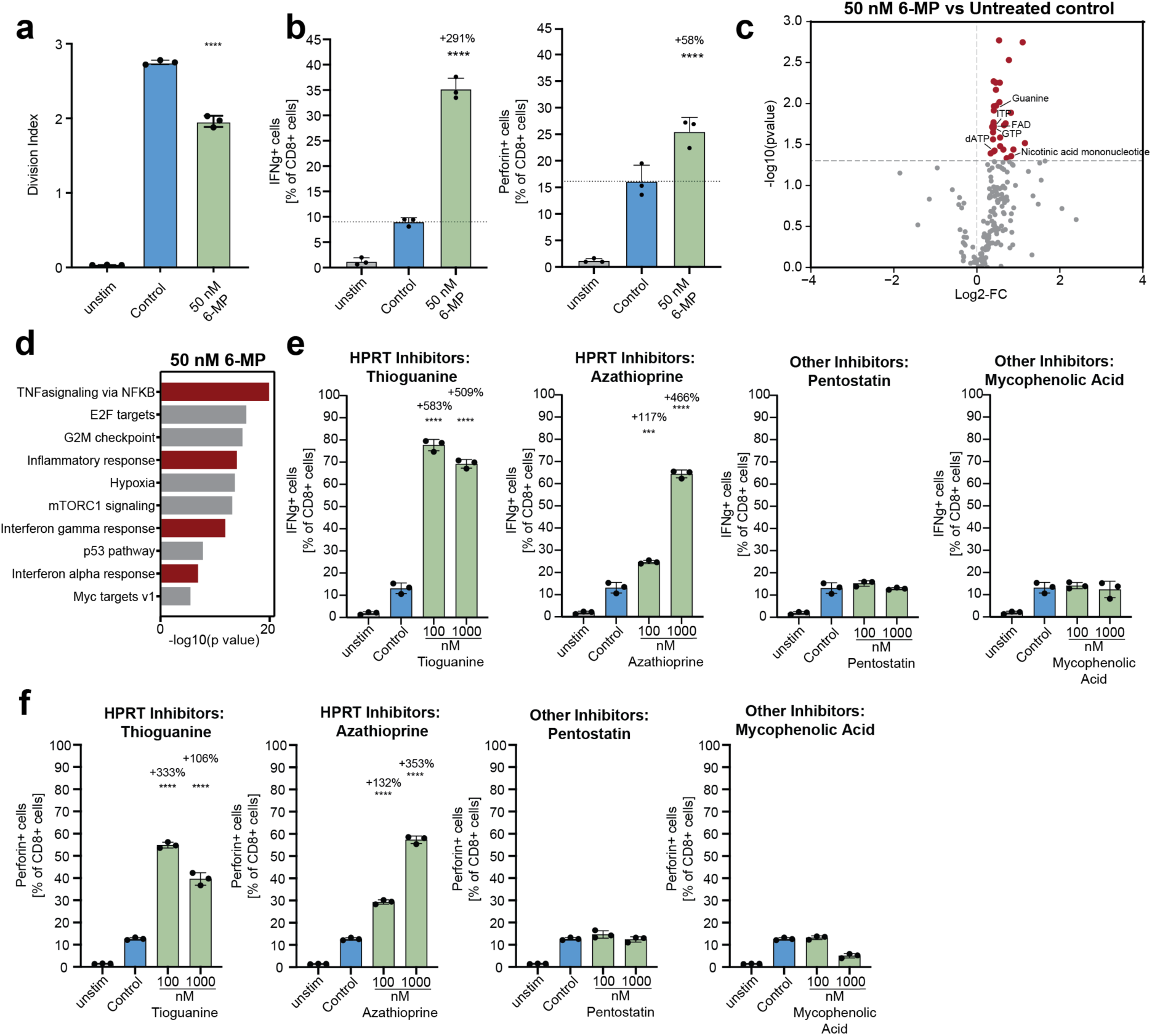
Pharmacological perturbation of the purine salvage pathway enhances CD8^+^ T cell effector function. a-b) Flow cytometry-based analysis of the effect of 6-mercaptopurine (abbreviation: 6-MP) on **a)** proliferative capacity and **b)** different parameters of effector function of CD8^+^ T cells stimulated *in vitro*. Statistical analysis: Independent t-test. The graph is a representative example of three independent experiments. **c)** Metabolomics analysis of the intracellular metabolite concentrations in CD8^+^ T cells after 6-mercaptopurine treatment compared to untreated control, evaluated 72 hours post stimulation *in vitro*. **d)** Gene set enrichment analysis of CD8^+^ T cells after 6-mercaptopurine treatment compared to untreated control, evaluated 72 hours post stimulation *in vitro*. **e-f)** FACS-based analysis of the effect of different inhibitors of purine metabolism on **e)** IFNγ expression and **f)** perforin expression in *in vitro*-stimulated CD8^+^ T cells. Data in all subfigures are presented as means ± SD. **a-f)** The sample size is n = 3.

We then assessed the specificity of 6-MP-mediated blockade of nucleotide synthesis on effector function by using mycophenolic acid and pentostatin, inhibitors targeting the enzymes inosine monophosphate dehydrogenase (IMPDH) and adenosine deaminase (ADA). These two compounds did not enhance CD8^+^ T cell effector function as determined by IFNγ and perforin expression (**Fig. 4e-f and Extended Data Fig. 4c-d**). Therefore, we concluded that the inhibition of nucleotide synthesis enhances the production of effector molecules in CD8^+^ T cells by reducing the metabolic demand for new purines produced via the resource-intensive purine *de novo* synthesis pathway. This reduction, in turn, spares biosynthetic and bioenergetic resources for effector function in a manner similar to that observed with purine supplementation.

To provide further evidence that the observed effects are not attributable to purinergic signaling, we employed inhibitors targeting cGAS-STING and purinergic signaling pathways. Contrary to the modulating effects observed with nucleosides such as adenosine and inosine, the purinergic signaling inhibitor ZM241385 failed to reverse the enhancement of CD8^+^ T cell effector function induced by 100 µM adenine (**Extended Data Fig. 4e**)^35^. Additionally, the cGAS-STING signaling inhibitors RU.521 and C-176 were also ineffective in reverting the enhanced effector function mediated by 6-mercaptopurine (**Extended Data Fig. 4f-g**). These findings further support the conclusion that the enhancement of CD8^+^ T cell effector function do not base on signaling activity.

### Slc43a3 facilitates transport of purine nucleobases during CD8^+^ T cell activation

To further delineate the mechanism of purine nucleobase uptake during the initiation and development of the antiviral CD8^+^ T cell response, we performed additional transcriptome analysis of LCMV-specific CD8+ T cells obtained from Doering *et al.*^31^. Interestingly, we unveiled a dynamic regulation of transporters facilitating transmembrane transport of metabolites in CD8^+^ T cells during viral infections including purine transporters (**Fig. 5a**). Notably, *Slc43a3*, which was recently identified as a purine nucleobase transporter, was upregulated in virus-specific CD8^+^ T cells during both infections, while expression levels of other transporters remained unchanged or decreased (**Fig. 5a**)^36^. Slc43a3 was found to selectively transport purine nucleobases across the membrane, but not nucleosides like adenosine which aligned with our observations on the effect of purine nucleobases and nucleosides on CD8^+^ T cells^36^. Based on these findings, we hypothesized that Slc43a3 mediates the uptake of purine nucleobases from the environment. To test this, we treated CD8^+^ T cells with the Slc43a3 inhibitor Decynium-22 (DC-22) and found that increasing concentrations reverted the effect of 100 µM adenine on enhanced effector function (**Fig. 5b and Extended Data Fig. 5a**)^36^. Moreover, metabolomics analysis of cells treated with 100 µM adenine, 10 nM DC-22 or a combination of both showed that 10 nM DC-22 successfully reverted the metabolic changes induced by the exogenous supplementation of adenine (**Fig. 5c**). Together, these findings led us to the conclusion that Slc43a3 facilitates the uptake of extracellular purine nucleobases in CD8^+^ T cells.

**Figure 5:**
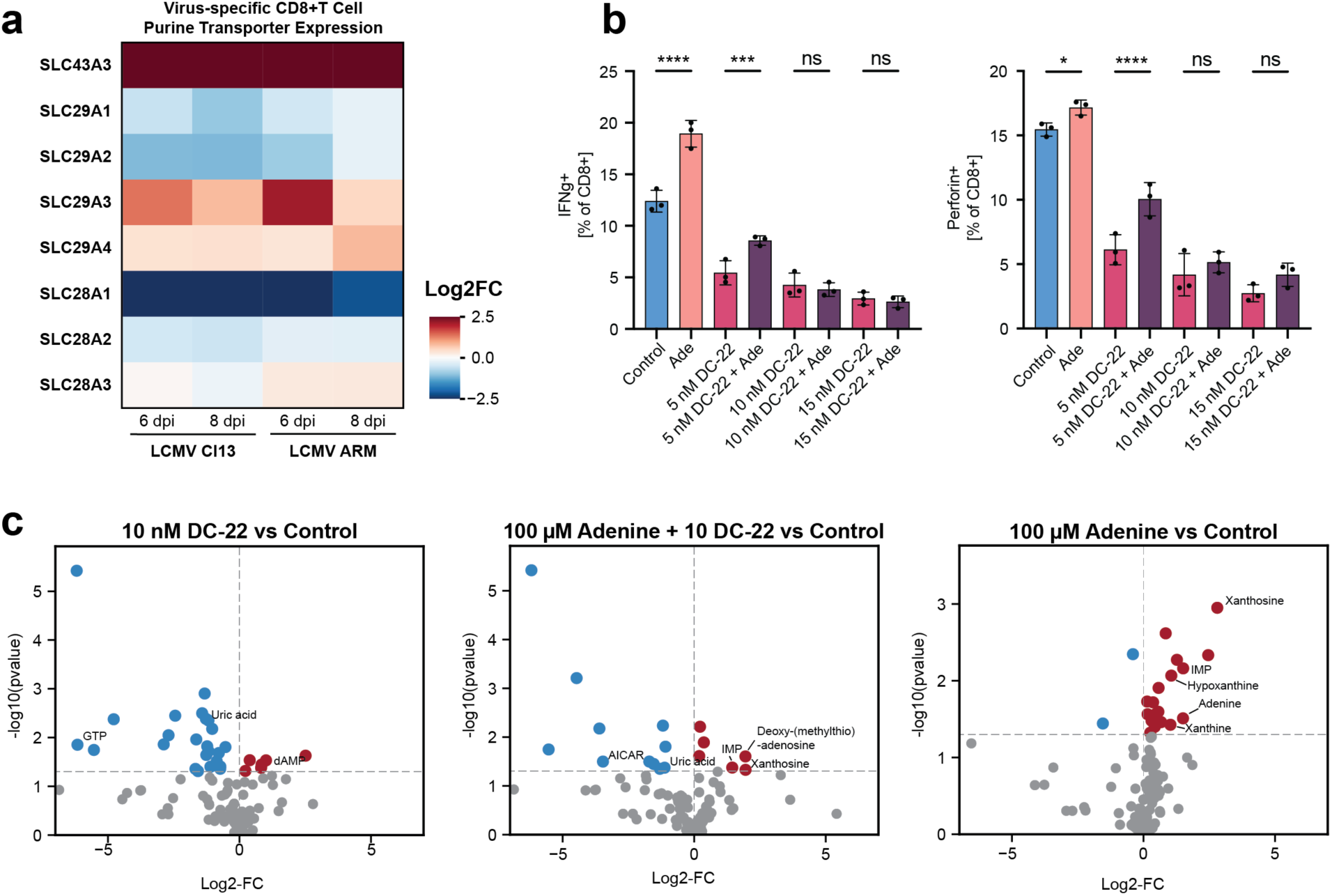
SLC43A3 facilitates transport of purine nucleobases during T cell activation. **a)** Differential gene expression analysis for purine transmembrane transporters in virus specific CD8^+^ T cells during LCMV ARM and LCMV Cl13 infection. Data obtained from Doering *et al*.^31^. **b)** Flow cytometry-based analysis of the effector function of *in vitro*-stimulated CD8^+^ T cells at different concentrations of decynium-22 (DC-22) with or without 100 µM adenine supplement. The graph is a representative example of two independent experiments. The data are represented as means ± SD with sample size n= 3. **c)** Metabolomic profiling of *in vitro*-stimulated CD8^+^ T cells treated with 10 nM DC-22, 100 µM Adenine or both compared to untreated control. The sample size is n= 3. Abbreviations: Ade – 100 µM adenine, DC-22 – Decynium-22.

### Endogenous elevation of serum hypoxanthine levels via allopurinol treatment enhances antiviral T cell responses

Based on our discovery of purine nucleobases as novel immunometabolites for the modulation of CD8^+^ T cell responses *in vivo*, we aimed to elucidate the effect of increased systemic levels of purine nucleobases by administering a purine-rich diet of 0.1% and 0.2% adenine to mice as reported previously^37^. However, even moderate adenine-enriched diets were not well tolerated in mice and side effects from the dietary treatment diminished potential positive effects from elevated serum purine levels (data not shown).

Hence, we resorted to elevate serum purine nucleobase levels via the endogenous route using allopurinol, an inhibitor of xanthine dehydrogenase (XDH) widely used in the clinics to prevent the conversion of purine nucleobases into xanthine and uric acid (**Fig. 3c**)^38^. First, we assessed the direct impact of allopurinol on CD8+ T cell effector function and proliferation *in vitro*. Our findings indicate that allopurinol does not affect these parameters across a broad dosage spectrum, up to 10 µM. (**Extended Data Fig. 6a,b**). Second, we verified that inhibition of XDH increases serum purine nucleobase levels (**Fig. 6a**). Leveraging this metabolic shift, we treated LCMV Cl13-infected mice with allopurinol at 4 dpi to augment their antiviral CD8^+^ T cell responses. The populations of GP33^+^ tetramer and NP396^+^ tetramer virus-specific antiviral CD8^+^ T cells were unaffected based on comparing percentages of epitope-specific CD8^+^ T cells (**Fig. 6b,c**). However, subsequent analysis of restimulated splenocytes and peripheral blood T cells revealed an enhancement in effector molecule production among CD8^+^ T cells (**Fig. 6d and Extended Data Fig. 6c**), accompanied by reduced viral loads in spleens and livers (**Fig. 6e**). Notably, serum levels of ALT (a liver-specific tissue damage marker) showed a slight decline (**Fig. 6f**). The cellular composition in terms of CD3^+^, CD4^+^, and CD8^+^ T cells, as well as T cell subsets, remained largely unaltered (**Extended Data Fig. 6d,e**). However, there was a significant increase in the central memory compartment among CD8^+^ T cells in both blood and spleen in the allopurinol-treated group (**Extended Data Fig. 6d,e**). In summary, these *in vivo* findings corroborate our observations from *ex vivo* experiments by demonstrating that elevation of purine nucleobase levels increases the effector function of antiviral CD8^+^ T cells without affecting proliferation.

**Figure 6:**
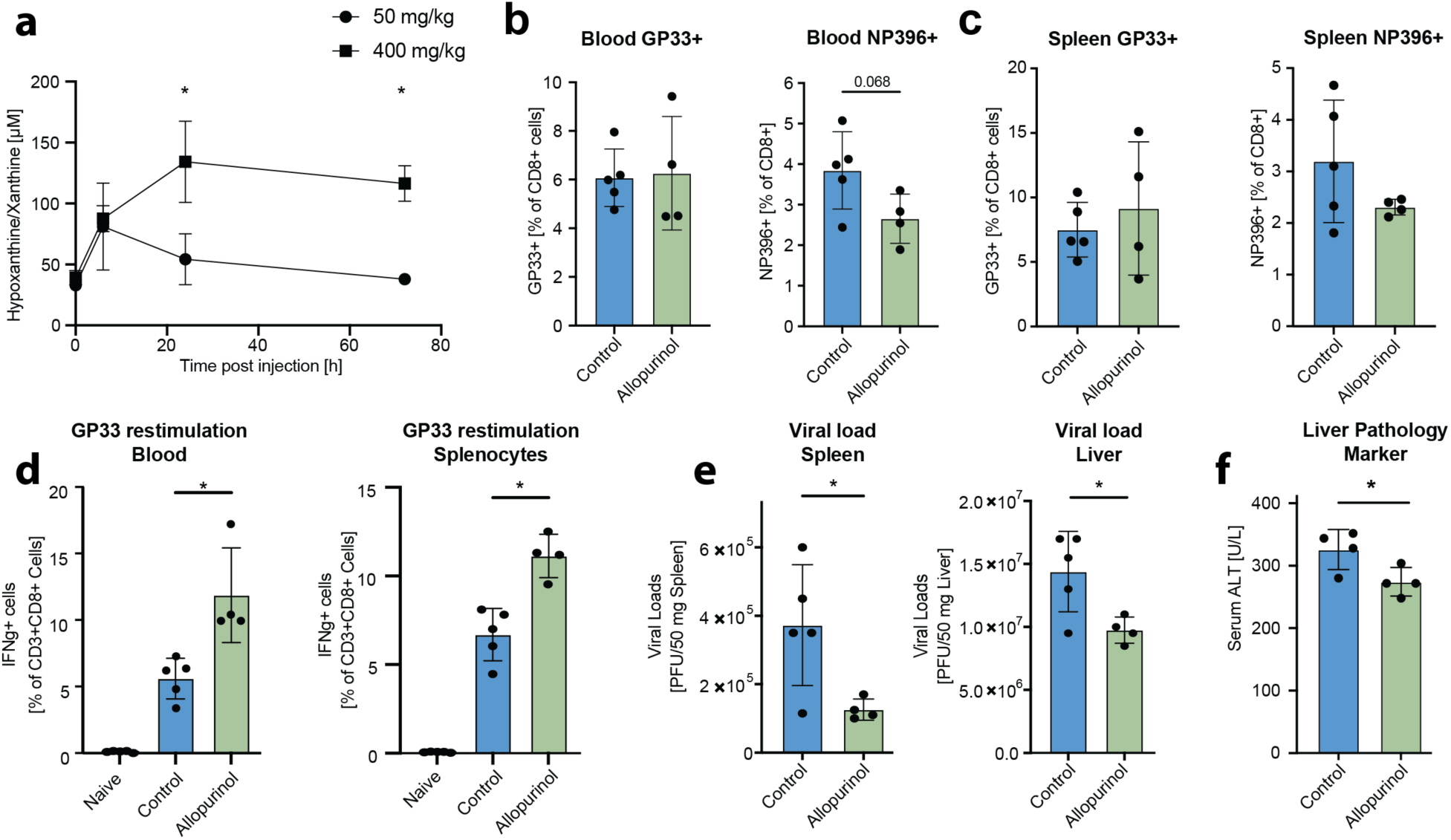
Elevation of serum purine nucleobase concentrations improves antiviral immune responses. **a)** Time-course analysis (longitudinal measurements) of serum hypoxanthine/xanthine concentration upon intraperitoneal administration of different doses of allopurinol. Statistical analysis: One-way ANOVA on longitudinal measurements with sample size is n=4 in 50 mg/kg treatment group and n=5 in 400 mg/kg treatment group. **b-c)** Flow cytometric analysis of GP33-and NP396-specific CD8^+^ T cells in **b)** blood and **c)** spleen. **d)** Flow cytometric analysis of peptide restimulated CD8^+^ T cells at 8 days post LCMV Cl13 infection. **e)** Viral loads in spleen and liver at 8 days post LCMV Cl13 infection. **f)** Serum ALT levels measured 8 days post LCMV Cl13 infection with and without allopurinol treatment. Statistical analysis: Independent t-test. The graph is a representative example of two independent experiments. **b-f)** Statistical analysis: Independent t-test. The graph is a representative example of two independent experiments. Sample size in the shown experiment n=5 in naïve and control or n=4 in treatment group, respectively. Data in all subfigures are presented as means ± SD

## Discussion

In this study, we investigated the interplay between metabolism and immunity in the context of viral infections and elucidated infection-associated modulation of systemic metabolism. Simultaneously, we harnessed viral infection models to identify novel immunometabolites and investigated the corresponding metabolic pathways that significantly shape CD8^+^ T cell responses. Our findings establish purine nucleobases as novel immunometabolites that are differentially regulated during viral infection and can be leveraged directly as metabolic fuels for CD8^+^ T cells or via pharmacological modulation of purine metabolism to enhance cytotoxic effector functions.

We provide a comprehensive metabolomics resource to explore the extensive spectrum of metabolic alterations associated with viral infections and their counteracting immune responses. Our repository encompasses a variety of murine and human viral infections, covering multiple RNA virus families including orthomyxoviruses, arenaviruses, and coronaviruses with different tissue tropisms and infection kinetics. Metabolomics analysis uncovered unique serum metabolic profiles associated with different stages and types of viral infections and their counteracting immune responses, demonstrating convergence or divergence in characteristic metabolic pathways as the infection progresses. These insights not only deepen our understanding of host-pathogen interactions but also provide a foundation for targeted interventions.

Two of the main findings emerging from our metabolomics data from various viral infection were that metabolic changes are more pronounced during the peak of the adaptive rather than the innate immune response and that purine metabolism consistently emerged as one of the most differentially regulated metabolic pathways in the analyzed murine infection models as well as in clinical data of human infections. These results highlight purine metabolism as a central immunometabolic hub during antiviral responses across species and types of viruses.

We further established a metabolite screen setup to identify novel immunometabolites among the differentially regulated serum metabolites observed. This targeted metabolite screening successfully yielded key metabolites known to be essential for T cell function, such as arginine and glutamine, thus validating the robustness of our findings^10,39^. We also identified purine nucleobases as new immunometabolites through this screen. Intriguingly, these compounds enhanced the effector function of CD8^+^ T cells without affecting their proliferative capacity. Previous research has demonstrated the critical role of purine *de novo* synthesis in maintaining nucleotide pools in proliferating T cells^9^. Here, we demonstrate that T cells can adapt metabolically by shifting from the resource-intensive purine *de novo* synthesis to the more efficient purine salvage pathway via uptake of purine nucleobases. This switch saves biosynthetic and bioenergetic metabolites, which may be reallocated to enhance the translation of effector molecules. We speculate that this metabolic flexibility not only allows CD8^+^ T cells to manage energy resources in varying metabolic environments but also enables them to overcome limiting conditions that may lack the biosynthetic substrates necessary for purine *de novo* synthesis, such as serine during LCMV Cl13 and MHV infections (**Fig. 1a**).

We conducted a set of control experiments to carefully validate metabolic utilization of purine nucleobases and to distinguish our findings from the domain of purinergic signaling.^15,40^ Purinergic signaling plays an important role in the regulation of immune cell activity within the microenvironment of inflamed tissues. It is facilitated by the release of nucleosides and nucleotides at very high concentrations, in the range of 10 mM, from necrotic cells.^15,40^. Hence, purine nucleotides and nucleosides, not nucleobases, have been extensively characterized as signaling molecules exerting immunomodulatory function^15,40^. Recent studies also showed that T cells could utilize the purine nucleoside inosine as an alternative carbon source to maintain proliferative capacity and that microbiome-derived inosine enhanced the effect of pro-inflammatory stimuli and immunotherapy^41,42^. However, the effects we described here are limited to purine nucleobases that were not reported before to have any purinergic signaling activity in immune cells. Our control experiments, including the use of inhibitors for classic purinergic signaling receptors like the A2A receptor and even the inhibition of the cGAS-STING pathway, suggested no signaling function underlying the effects of purine nucleobases that we observed. Instead, metabolite tracing data demonstrated the conversion of imported purine nucleobases to nucleotides via the purine salvage pathway. We, thus, conclude that purine nucleobase uptake substitutes purine *de novo* synthesis and thereby frees up cellular biosynthetic and bioenergetic resources which contributes to enhanced effector function. This mechanism offers new insights into the complex and multifaceted role of purine metabolism as an immunometabolic pathway.

The application of purine analogs unveiled further critical insights into this immunometabolic hub for the allocation of biosynthetic and bioenergetic resources within T cells, particularly between proliferation and the production of effector molecules. Drugs targeting nucleotide synthesis are commonly used in the clinics to block the establishment of T cell responses via suppression of proliferation. Surprisingly, our study found that compounds like 6-mercaptopurine and azathioprine, specifically at lower doses, boosted CD8^+^ T cell effector function with minor inhibition of cell proliferation. Hence, our results showed that these compounds can effectively redirect cellular resources from proliferation to effector function. Together, the experiments with purine nucleobases and purine analogs demonstrated not only the stringent regulation of cellular bioenergetic and biosynthetic resource allocation but also the remarkable adaptability of CD8^+^ T cells in managing these metabolic resources for proliferation and effector function.

Purine nucleobases are abundant in various foods like meat, and the consumption of external purines depends largely on individual dietary lifestyle. As purine nucleobases appear to act as biosynthetic fuel for T cell responses, further research could explore how different diets – specifically purine-rich omnivorous versus low-purine plant-based diets – might influence antiviral immunity. In conclusion, our study demonstrates a novel regulatory role of purine metabolism in supporting CD8^+^ T cell responses and could open new avenues for immunometabolic targets for immunotherapy.

## Data availability

The data that support the findings of this study are available from the corresponding author (A.B.) upon request. This study did not generate new unique reagents. Metabolomics source data used to generate Fig. 1a-d, Extended Data Fig. 1a-e, Fig. 3a,c and Fig. 4c are provided with this paper in the Extended Data Tables 1 and 3. Metabolite screen source data used to generate Fig. 2b and Extended Data Fig. 2a are provided with this paper in Extended Data Table 2. RNAseq source data used for GSEA in Fig. 2E and Fig. 3D are available at the European Nucleotide Archive under the accession number PRJEB70319.

## Material and Methods

### Mice

C57BL/6J mice, initially sourced from The Jackson Laboratory, were bred and housed under specific pathogen-free (SPF) conditions at the Institute for Molecular Biotechnology of the Austrian Academy of Sciences and the Animal Facility of the Medical University of Vienna in Vienna, Austria or directly purchased from Charles River. All animal experiments, conducted in individually ventilated cages, adhered to the approved license (BMWFW-2020-0.406.011) and ethical guidelines set by the institutional committees at the Medical University of Vienna’s Department for Biomedical Research. The mice, aged between 8 and 12 weeks old, were age-and sex-matched for each experiment. For studies involving murine hepatitis virus (MHV), the mice were kept under SPF conditions at the Kantonsspital St. Gallen Medical Research Center in Switzerland, with MHV experiments performed in compliance with the license (SG/14/18.30861) approved by the relevant federal and cantonal ethical committees.

### Viruses and cell lines

BHK-21 cells, derived from the kidneys of five newborn hamsters (ATCC CCL-10), were used to culture Lymphocytic choriomeningitis virus (LCMV). The virus’s concentration was determined through an adapted focus-forming assay utilizing Vero cells (ATCC CCL-81), which originate from the kidneys of female African green monkeys^43^. For the infection experiments, mice received an intravenous injection of 2×10 focus-forming units (FFU) of LCMV ARM or LCMV Cl13.

Murine hepatitis virus strain A59 (MHV) was propagated in 17CL1 cells, a spontaneously transformed cell line from BALB/c mouse embryos^44^. The viral titer for MHV was determined using a standard plaque assay on L929 cells, which are male mouse fibroblasts (ATCC CCL-1)^45^. Mice were administered an intraperitoneal injection of 10^3 plaque-forming units (PFU) of MHV.

### Infection and sample processing

Mice were euthanized at specified time points noted in the figure legends using cervical dislocation. Immediately after, their tissue samples were flash-frozen using liquid nitrogen and then preserved at-80°C for subsequent analysis. To obtain serum for metabolomics and blood chemistry analyses, blood samples were collected from the tail vein in tubes covered with clot activator (Microvette® 100 Z, Sarstedt AG, 20.1280.100) at the designated times prior to euthanasia, followed by centrifugation at 10,000 rpm for 5 minutes at 4°C. The collected serum was then placed in fresh tubes and kept at-80°C for future examinations.

### Serum metabolomics of hepatitis C virus (HCV) patients

Serum samples were obtained from 16 patients (males; mean age: 41.1± 7.5 years) with HCV infection prior to antiviral HCV treatment and after sustained virologic response (i.e., assessed 3-6 months after treatment). HCV-RNA levels (viral loads) and serum ALT levels were determined to confirm acute infection or virus clearance, respectively. The clinical trial adhered to the Declaration of Helsinki and was approved by the local ethics committee of the Medical University of Vienna (MUV-EC Nr: 1527/2017)^46^.

### Pharmacological perturbation

Allopurinol (Sigma Aldrich) was administered to mice via intraperitoneal injection at a dosage of 50 and 400 mg/kg in 10% Tween-80/PBS as described elsewhere^47^. Mice were closely monitored for any indications of distress or adverse effects throughout the study. Treatment was started four days after LCMV infection. Control mice received the same volume of vehicle (10% Tween-80/PBS).

### T Cell isolation

Murine CD8^+^ T cells were isolated from splenocytes using the MojoSort Mouse CD8 T Cell isolation kit (480035, BioLegend) according to the manufacturer’s protocol in combination with magnetic columns for column-based negative selection.

### *In vitro* CD8^+^ T cell stimulation and metabolite screen

Murine splenic CD8^+^ T cells were seeded in 384-well plates at a density of 15,000 cells per well. For metabolic and substrate utilization studies, cells were cultivated in RPMI enriched with metabolites of interest and supplemented with 10% dialysed FBS (F0392, Sigma Aldrich), 1% Penicillin/Streptomycin, 55 µM β-Mercaptoethanol, 50 U/mL recombinant murine IL-2 (212-12, Peprotech) and Dynabeads Mouse T-Activator CD3/CD28 (11456D, Thermo Fischer Scientific) at a 1:1 bead to cell ratio. For studying amino acid utilization, synthetic amino acid-free RPMI was selectively enriched or depleted of specific amino acids. After 72 hours incubation at 37 °C and 5 % CO_2_, cells were restimulated with PMA/Ionomycin and brefeldin A and monensin (00-4975, Thermo Fisher Scientific). After 6 hours restimulation, cells were fixated with 0.5 % PFA and 0.15% Triton X100 and stained with DAPI, anti-IFNγ-PE (1:250 dilution) and anti-CD44-AF488 (1:400 dilution) over night at 4 °C and 6 hours at room temperature before washing and imaged immediately afterwards. Imaging readouts were conducted with the PerkinElmer Opera Phenix High-Content High-Throughput Imaging System at 20x resolution acquiring nine fields per well for all three channels with the following emission filters (DAPI: 435-480nm, PE: 570-630 nm, AF488: 500-550 nm).

### *In vitro* CD8^+^ T cell stimulation and flow cytometry

Plates for CD8^+^ T cell stimulation were treated with antibody solutions of murine aCD3 (1 µg/mL, 553238, BD Biosciences) and murine aCD28 (2.5 µg/mL, 553295, BD Biosciences) antibodies in PBS over night at 4 °C. 96-well plates were treated with 100 µL, 48-well plates were treated with 250 µL per well. Murine splenic CD8^+^ T cells were seeded in 48-well plates at a density of 400,000 cells per well or in 96-well plates at a density of 50,000 cells per well. For metabolic and substrate utilization studies, cells were cultivated for 72 hours at 37 °C and 5% CO_2_ in RPMI supplemented with metabolites of interest and 10% dialyzed FBS (F0392, Sigma Aldrich), 1% Penicillin/Streptomycin, 55 µM β-Mercaptoethanol (BME, Sigma Aldrich) and 50 U/mL recombinant murine IL-2 (212-12, Peprotech).

For proliferation studies, naïve cells were stained before plating using Cell Proliferation Dye eFluor 450 (ThermoFisher Scientific, 65-0842-85) according to the manufacturer’s protocol. Proliferation readouts are reported as a division index of viable cells, describing the average number of cell divisions per cell including undivided cells.

For tetramer staining, cells were first suspended in a 25 μL solution of PBS with GP33 (dilution 1:500) and NP396 (dilution 1:250) tetramers from the NIH Tetramer Core Facility, followed by a 15-minute incubation at 37°C. Subsequently, 25 μL of PBS mixed with anti-CD16/32 (Biolegend; dilution 1:200) was added, and the mixture was incubated at room temperature for 10 minutes. Then, a 25 μL master mix containing a selection of surface marker antibodies (including anti-CD8.2b, anti-CD3 and others, all from Biolegend at a 1:200 dilution in PBS) along with Fixable Viability Dye eFluor 780 (eBioscience; 1:2000 in PBS) was added for a 20-minute incubation at 4°C. After washing with FACS buffer (PBS with 2% FCS), cells were fixed in 4% Paraformaldehyde (Sigma) in PBS for 10 minutes, washed twice more with FACS buffer, resuspended in 100 μL, and analyzed via flow cytometry.

For intracellular cytokine staining (ICS), cell pellets were reconstituted in 50 μL of RPMI 1640 medium (GIBCO) enriched with 10% FCS (PAA) and 1% Penicillin-Streptomycin-Glutamine (Thermo Fisher Scientific), plus 55 μM β-mercaptoethanol (Sigma). This medium also included LCMV peptides (1:1000, Peptide 2.0 Inc.) and a Protein Transport Inhibitor Cocktail (eBioscience, #00-4980-03; 1:500, Thermo Fisher Scientific). Cells were treated with a Cell Stimulation Cocktail (eBioscience, #00-4970-93) as a positive control and incubated for 4 hours at 37°C. Surface antigens were stained as previously described. Following this, a 25 μL master mix of selected antibodies in FACS buffer with 0.05% saponin (Sigma, 47036) for intracellular targets (including anti-IFNγ, anti-Perforin, anti-Gzmb; all from Biolegend, all at 1:200 dilution) was added and incubated for 90 minutes at 4°C. Finally, the cells were washed twice with FACS buffer, resuspended in 100 μL, and subjected to flow cytometric analysis.

### ELISA for effector molecule quantification

IFNγ and TNFα were quantified in 100 µL cell media supernatants after 72 hours incubation of *in vitro*-stimulated murine splenic CD8^+^ T cells according to the manufacturer’s protocols (IFNγ: Thermo Fisher Scientific, #88-7314-88; TNFα: Thermo Fisher Scientific, #88-7324-76). Signals were recorded in the wavelength substraction mode (570 nm signal substracted from 450 nm) with a SpectraMax i3x Multi-Mode Microplate Reader in 96-well plate format. Effector molecule concentrations were determined by comparing signals from samples and provided standards and normalization by cell count.

### Metabolomics analysis

Mouse serum, obtained as previously described, was extracted by adding ice-cold methanol and cleared extracts were dried under nitrogen. Samples were taken up in MS-grade water and mixed with the heavy isotope labelled internal standard mix. A 1290 Infinity II UHPLC system (Agilent Technologies) coupled with a 6470 triple quadrupole mass spectrometer (Agilent Technologies) was used for the LC-MS/MS analysis. The chromatographic separation for samples was carried out on a ZORBAX RRHD Extend-C18, 2.1 x 150 mm, 1.8 um analytical column (Agilent Technologies). The column was maintained at a temperature of 40°C and 4 µL of sample was injected per run. The mobile phase A was 3% methanol (v/v), 10 mM tributylamine, 15 mM acetic acid in water and mobile phase B was 10 mM tributylamine, 15 mM acetic acid in methanol. The gradient elution with a flow rate of 0.25 mL/min was performed for a total time of 24 min. Afterwards back-flushing of the column using a 6port/2-position divert valve was carried out for 8 min using acetonitrile, followed by 8 min of column equilibration with 100% mobile phase A. The triple quadrupole mass spectrometer was operated in negative electrospray ionization mode, spray voltage 2 kV, gas temperature 150 °C, gas flow 1.3 L/min, nebulizer 45 psi, sheath gas temperature 325 °C, sheath gas flow 12 L/min. The metabolites of interest were detected using a dynamic MRM mode. The MassHunter 10.0 software (Agilent Technologies) was used for data processing. Seven-point calibration curves with internal standardization was constructed for the quantification of metabolites. The log2 fold-changes for differential metabolite modulation were determined based on pairwise comparisons of metabolite concentrations between the replicates of uninfected control samples versus the replicates of infected samples.

### Metabolite tracing

CD8^+^ T cells were stimulated *in vitro* in cell culture media with dialyzed FBS (F0392, Sigma Aldrich) for 72 hours at 37°C with supplements of 8-^13^C heavy-isotope labelled adenine before measurement. To elucidate the metabolic utilization of ^13^C-labelled adenine (8-^13^C, 95%, Cambridge Isotope Labs) the same extraction and LC-MS set-up as for the targeted metabolomics was used, except no heavy isotope-labelled internal standards were added, an adapted dynamic MRM list focused on purines, pyrimidines, and related control metabolites were constructed monitoring both light and heavy metabolites, and signals were quantified as area under the curve (AUC) without absolute quantification. Ratios of AUC between heavy and light metabolites were compared for normalization.

### Measurement of intracellular ATP levels using CellTiter-Glo^®^

Intracellular ATP content in CD8+ T cells was assessed with CellTiter-Glo^®^ Luminescent Cell Viability Assay (Promega) after 72 h incubation at 37 °C with or without purine nucleobase supplements. Briefly, CD8^+^ T cells were resuspended in RPMI 1640 and seeded at a density of 200,000 cells per well in 96-well plates (Corning 3764) and measurements conducted according to the manufacturer’s instructions.

### Measurement of translation rates using SCENITH™

Translation rates were determined using the SCENITH™ method using the SCENITH kit reagents (www.scenith.com) by detecting puromycin incorporation into proteins in CD8^+^ T cells for 30 minutes after 72 h incubation at 37 °C with or without additional treatments as described before^34^. For the measurement, cells were resuspended in RPMI 1640 supplemented with 10% dialysed FBS and 10 µg/mL Puromycin dihydrochloride (HY-B1743A, MedChemExpress) and incubated for 30 minutes at 37 °C. Cells were then stained for flow cytometry-based analysis as previously described, and stained with AF647-conjugated anti-puromycin antibody supplied by Dr. Argüello to assess puromycin incorporation.

### Sample collection for RNA sequencing

48-well plates for the creation of RNA sequencing samples were treated with antibody solutions of murine aCD3 and murine aCD28 as described for the Flow cytometry setup before. Murine splenic CD8^+^ T cells were seeded in 48-well plates at a density of 400,000 cells per well and harvested after 72 hours.

### NGS Library Preparation for RNA sequencing

The amount of total RNA was quantified using the Qubit 2.0 Fluorometric Quantitation system (Thermo Fisher Scientific, Waltham, MA, USA) and the RNA integrity number (RIN) was determined using the Experion Automated Electrophoresis System (Bio-Rad, Hercules, CA, USA). RNA-seq libraries were prepared with the TruSeq Stranded mRNA LT sample preparation kit (Illumina, San Diego, CA, USA) using Sciclone and Zephyr liquid handling workstations (PerkinElmer, Waltham, MA, USA) for pre-and post-PCR steps, respectively. Library concentrations were quantified with the Qubit 2.0 Fluorometric Quantitation system (Life Technologies, Carlsbad, CA, USA) and the size distribution was assessed using the Experion Automated Electrophoresis System (Bio-Rad, Hercules, CA, USA). For sequencing, samples were diluted and pooled into NGS libraries in equimolar amounts.

### Next-Generation Sequencing and Raw Data Acquisition for RNA sequencing

Expression profiling libraries were sequenced on HiSeq 3000/4000 instruments (Illumina, San Diego, CA, USA) following a 50-base-pair, single-end recipe. Raw data acquisition (HiSeq Control Software, HCS, HD 3.4.0.38) and base calling (Real-Time Analysis Software, RTA, 2.7.7) was performed on-instrument, while the subsequent raw data processing off the instruments involved two custom programs based on Picard tools (2.19.2). In a first step, base calls were converted into lane-specific, multiplexed, unaligned BAM files suitable for long-term archival (IlluminaBasecallsToMultiplexSam, 2.19.2-CeMM). In a second step, archive BAM files were demultiplexed into sample-specific, unaligned BAM files (IlluminaSamDemux, 2.19.2-CeMM).

### Blood chemistry analysis

Blood for the assessment of alanine aminotransferase (ALT) and aspartate aminotransferase (AST) levels was collected in MiniCollect EDTA tubes (Greiner Bio-One). Serum was separated by spinning at 4,000 rpm for 10 minutes at 4°C. Serum was diluted in a 1:8 ratio with PBS. Samples were kept chilled, shielded from light, and sealed until they were analyzed. ALT and AST levels were measured using a spectrophotometric method on a Roche Cobas C311 Analyzer.

### Serum xanthine/hypoxanthine measurements

Xanthine/hypoxanthine concentrations in mouse serum were determined using the Xanthine/hypoxanthine assay kit (Sigma-Aldrich, #MAK186) according to the manufactuerer’s protocol. Proteins were removed from serum samples using 3 kDa Amicon ultracentrifugation filters (Merck, UFC5003). Samples were prepared for fluorimetric assay measurement, signals recorded with SpectraMax i3x Multi-Mode Microplate Reader and concentrations of xanthine/hypoxanthine determined by comparine signals of samples to the standard curve.

## Data Processing and Statistical Analysis

### Metabolite set enrichment analysis

Metabolite set enrichment was conducted using metabolite concentration tables and the Quantitative Enrichment Analysis tool of MetaboAnalyst 5.0^48^. Metabolite concentration data were log-transformed and enriched metabolite sets were assessed based on the KEGG pathways. Only metabolite sets with q value of enrichment < 0.05 were considered.

### RNAseq data processing and gene set enrichment analysis

Sequencing data were converted to fastq files using bedtools^49^. Adaptor sequences and low-quality bases were removed using trimmomatic (0.38), with parameters set to 2:30:10 LEADING:3 TRAILING:3 SLIDINGWINDOW:4:15 MINLEN:30^50^. Transcript and gene abundance was deduced with Salmon (1.4.0) with correction for sequence and GC content bias^51^. Reference mouse transcriptome mm10 was deduced from refgenie (0.12.0) with refgenconf (0.12.2)^52^. Obtained reads per gene counts were imported into R and pairwise differential gene expression analysis was performed using the DESeq2 package^53^. The EnrichmentBrowser (2.20.7) package was used to examine the resulting gene list and the set of differentially expressed genes, defined by adjusted p-value <0.05, for overrepresented gene sets as as annotated by Molecular Signatures Database (msigdbr 7.4.1), using the overrepresentation analysis methods (ORA) implemented in the EnrichmentBrowser package^54^.

### Principal component analysis

Principal component analysis (PCA) was performed using Scikit-learn and standard-scaled metabolite concentration tables^55^. Ellipses around data points for each condition represent covariance confidence intervals with a radius of two standard deviations.

### Statistical information

Data are presented as arithmetic mean ± SD. Sample sizes are indicated in the figure legends. Longitudinal metabolomics measurements in mice during viral infections were analyzed with two-sided paired t-test (naïve vs 2 dpi or 8 dpi, respectively) with two-stage step-up Benjamini-Hochberg procedure to control the false discovery rate (FDR) (alpha = 0.1). Statistical significance in screens was evaluated using two-sided independent t-test with Benjamini-Hochberg procedure to control the FDR (alpha = 0.1). If not indicated differently, statistical significances in other experiments were calculated with one-way analysis of variance (ANOVA). Stars indicate significance levels based on p values as follows: ns = not significant, * - p < 0.05, ** - p < 0.01, *** - p < 0.001, **** - p < 0.0001.

## Acknowledgements

We want to thank Sarah Niggemeyer, Jenny Riede and Nicole Fleischmann for animal caretaking; Burkhard Ludewig for providing samples from infection experiments with MHV; Iciar Serrano Sanchez and Juan Sanchez Avila from the Molecular Discovery Platform at CeMM for metabolomics analysis and data processing; Thomas Penz, Michael Schuster, Martin Senekowitsch, Daniele Barreca, and Christoph Bock from the Biomedical Sequencing Facility at CeMM for RNA sequencing and initial data analysis; Clarissa Campbell for support and feedback. The following reagents were obtained through the NIH Tetramer Core Facility: LCMV MHCI tetramers GP33 (conjugated to Phycoerythrin [PE]) and NP396 (conjugated to Allophycocyanin [APC]). This project was funded with support of the European Research Council (ERC) under the European Union’s Horizon 2020 research and innovation program (grant agreement no. 677006, “CMIL” to A. Bergthaler). A DOC Fellowship from the Austrian Academy of Sciences supported J.-W. G. and A. L. RJA was supported by the grant ANR PRC MetaNiche N° ANR-22-CE15-0015-02.

---

**Extended Data Figure 1:**
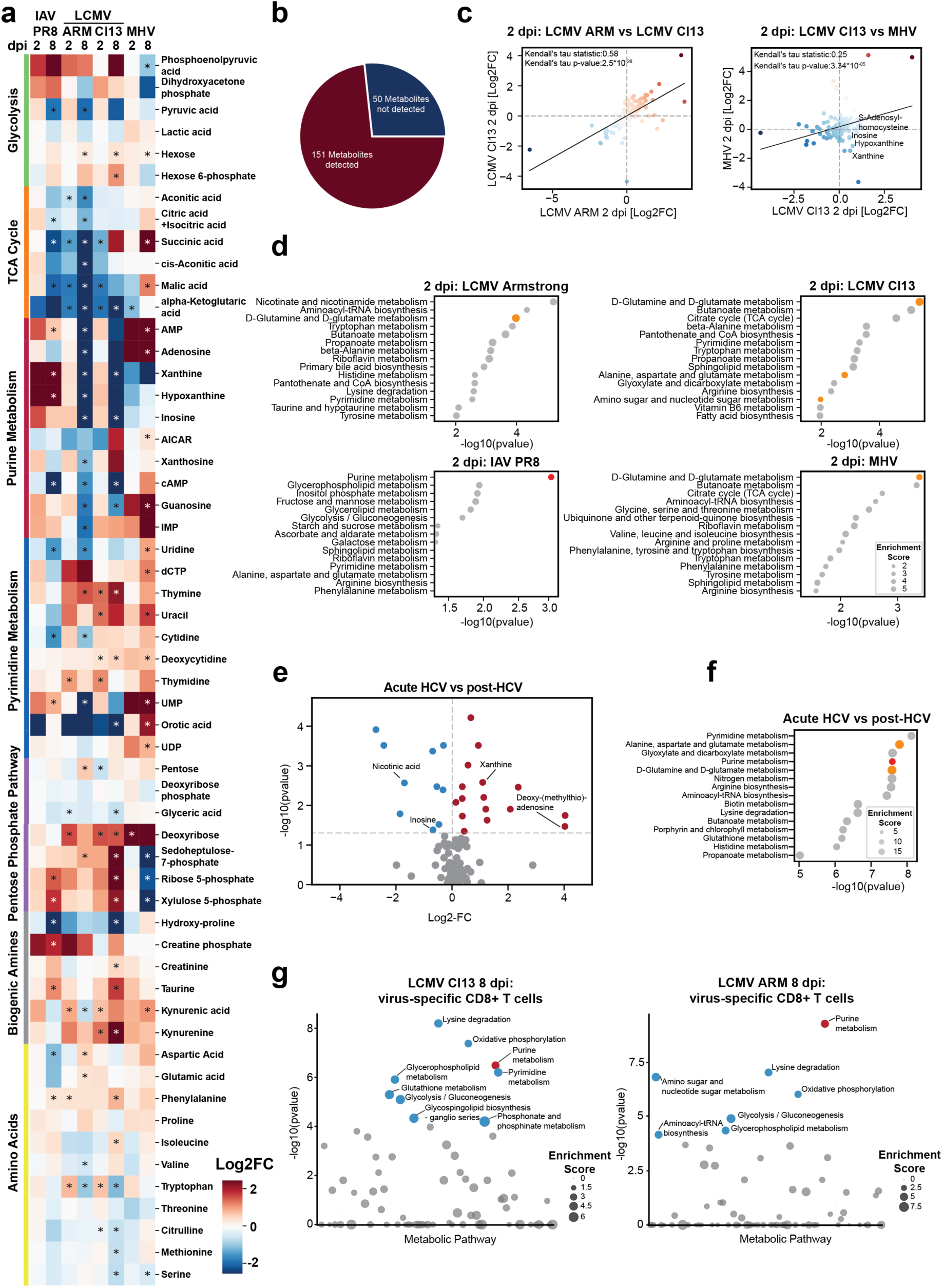
Comprehensive analysis of systemic metabolism reveals dynamic patterns in serum metabolite concentrations during viral infections. **a)** Longitudinal profiling of serum metabolite changes in five age-and sex-matched mice per group infected with LCMV Armstrong (LCMV ARM), LCMV Cl13, Influenza A virus Puerto Rico 8 (IAV PR8) or murine hepatitis virus (MHV). Longitudinal metabolomics measurements were analyzed with paired t-tests (naïve vs 2 dpi or 8 dpi, respectively) with two-stage step-up Benjamini-Hochberg procedure to control the FDR (alpha = 0.1). **b)** Number of metabolites of the panel detected in at least 10 samples. **c)** Correlation matrix of serum metabolite changes 2 dpi. **d)** Metabolite set enrichment analysis indicating perturbed metabolic pathways during infection with LCMV ARM, LCMV Cl13, IAV PR8, and MHV at 2 dpi. Purine metabolism is emphasized in red, with associated metabolic pathways marked in yellow. **e)** Metabolic changes in serum during acute HCV infection compared to cured patients post treatment. Statistical Analysis: Wilcoxon Rank-sum Test. **f)** Metabolite set enrichment analysis indicating perturbed metabolic pathways during acute HCV infection. **f)** Gene set enrichment analysis of metabolic pathway genes (according to KEGG annotation) in virus-specific T cells during LCMV Cl13 and LCMV Armstrong infection at 8 dpi. Data obtained from Doering et al.^31^

**Extended Data Figure 2:**
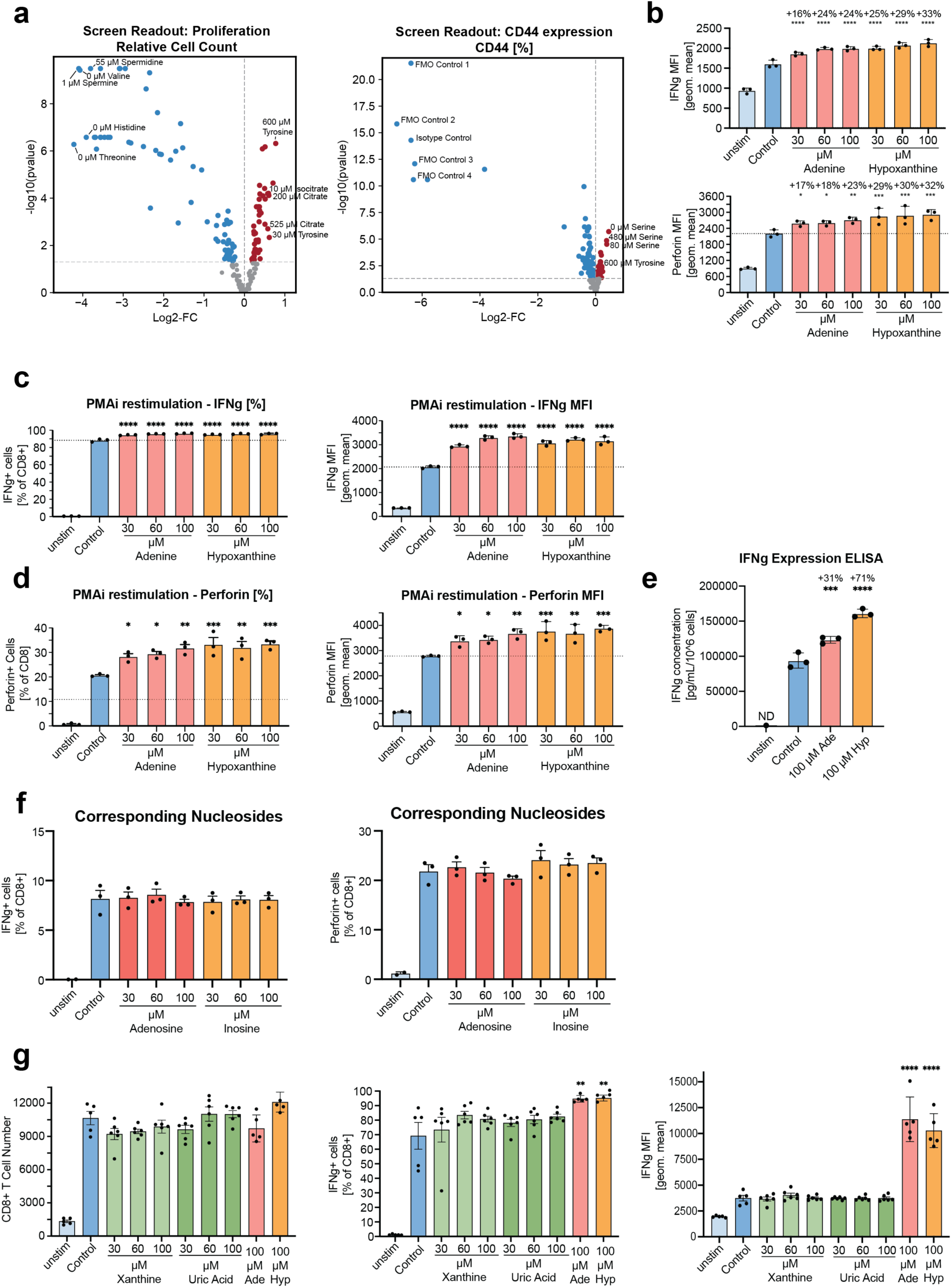
a) Results of proliferation readout and CD44 expression readout in high-throughput metabolite screen setup with sample size n = 6. **b)** Flow cytometry-based validation of metabolite screen outcomes examining the impact of purine nucleobases on mean fluorescence intensity (MFI) of IFNγ-and Perforin-producing CD8^+^ T cells. The graph is a representative example of two independent experiments. **c-d)** Proportion and MFI intensity of **c)** IFNγ-producing and **d)** perforin-producing CD8^+^ T cells upon treatment with purine nucleobase supplements and restimulation with phorbol myristic acetate and ionomycin (PMAi) *in vitro*. The graph is a representative example of two independent experiments. **e)** ELISA-based validation of metabolite screen outcomes examining the impact of purine nucleobases on effector function. The graph is a representative example of two independent experiments **f)** High-throughput image-based analysis of the effect of xanthine and uric acid on proliferation and effector function of CD8^+^ T cells *in vitro*. **g)** Flow cytometry-based analysis of the effect of the nucleosides adenosine and inosine on the effector function of CD8^+^ T cells *in vitro*. The graph is a representative example of three independent experiments. Data in all subfigures are presented as mean ± SD. c-f) Sample size in flow cytometry readout is n = 3. g) Sample size in image-based screen is n = 5 in control group and n = 6 in treatment and unstimulated groups. Abbreviations: Ade - 100 µM adenine; Hyp – 100 µM hypoxanthine

**Extended Data Figure 3:**
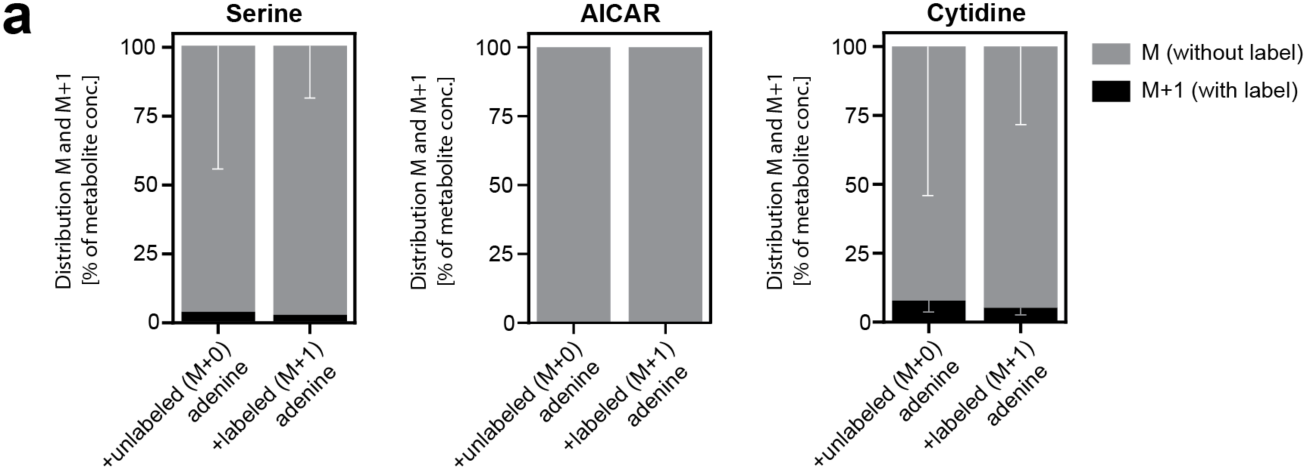
Salvaging purine nucleobases spares biosynthetic and bioenergetic resources in CD8^+^ T cells for enhanced effector function. **a)** Mass spectrometry-based analysis of the distribution of heavy isotope-labelled C atoms in cellular metabolites. CD8^+^ T cells were stimulated *in vitro* in cell culture media with dialyzed FBS and supplemented with 100 µM 8-^13^C heavy-isotope labelled adenine for 72 hours before measurement. Sample size is n = 3. Data are represented as percentage of total with standard deviation.

**Extended Data Figure 4:**
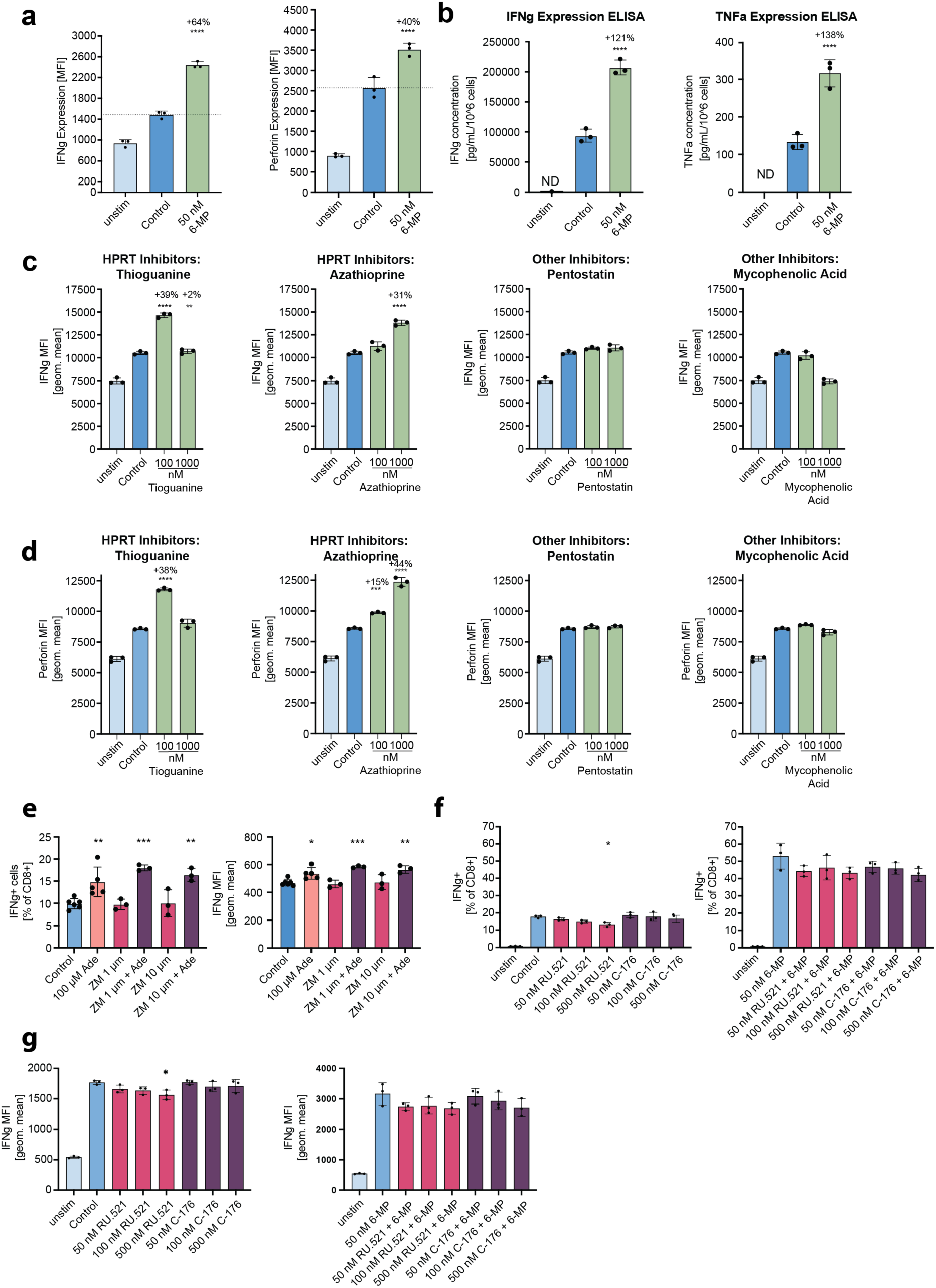
Pharmacological perturbation of purine salvage pathway enhances CD8^+^ T cell effector function. **a)** Flow cytometry-based validation of metabolite screen outcomes examining the impact of purine nucleobases on effector function. **b)** ELISA-based analysis of the effect of 6-mercaptopurine on effector function. The graph is a representative example of three independent experiments. **c-d)** FACS-based analysis of the effect of different inhibitors of purine metabolism on **c)** IFNγ expression and **d)** perforin expression in *in vitro*-stimulated CD8^+^ T cells. The graph is a representative example of tree independent experiments. **e)** Flow cytometry-based assessment of the efficacy of the purinergic signaling inhibitor ZM241385 to revert the effects of 100 µM adenine treatment. **f-g)** Flow cytometry-based analysis of the efficacy of cGAS-STING signaling inhibitors in inhibiting the effect of 6-mercaptopurine on **f)** proportion and **g)** mean fluorescence intensity (MFI) of IFNg-producing CD8^+^ T cells *in vitro* 72 hours post stimulation. Data in all subfigures are presented as mean ± SD with sample size a-d,f-g) n = 3 or e) n = 5 in control and Ade treatment and n = 3 in inhibitor treatments. Abbreviations: Ade - 100 µM adenine; 6-MP – 50 nM 6-mercaptopurine

**Extended Data Figure 5:**
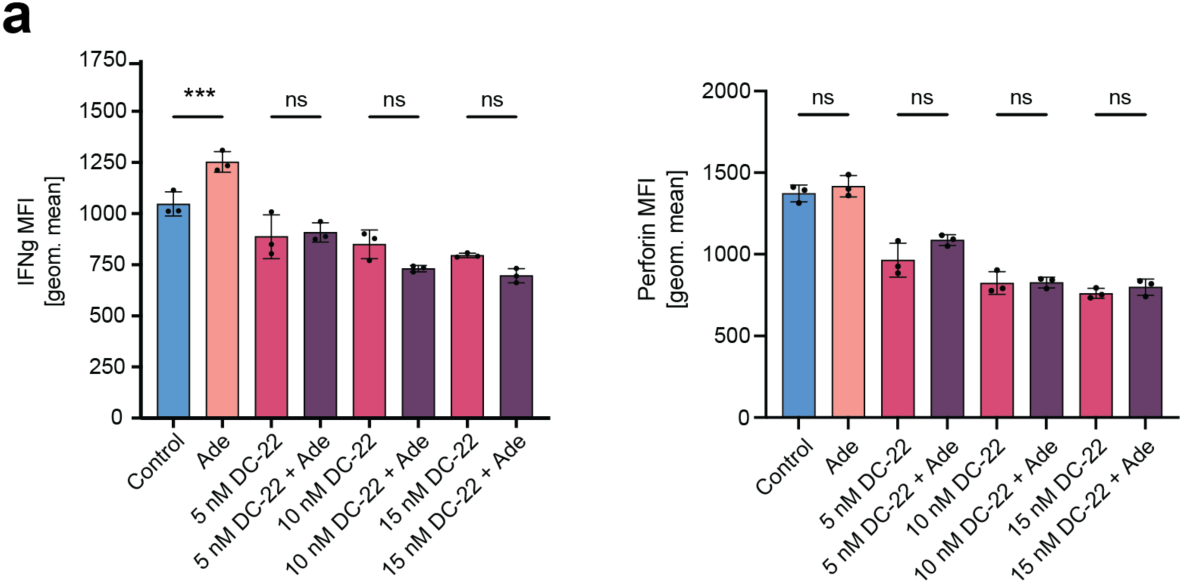
SLC43A3 facilitates purine nucleobases transport during T cell activation. **a)** Flow cytometry-based analysis of different parameters of the effector function of *in vitro*-stimulated CD8^+^ T cells at different concentrations of decynium-22 (DC-22) with or without 100 µM adenine supplement. The graph is a representative example of two independent experiments. Data in all subfigures are presented as mean ± SD with sample size n = 3. Abbreviations: Ade - 100 µM Adenine, DC-22 – Decynium-22.

**Extended Data Figure 6:**
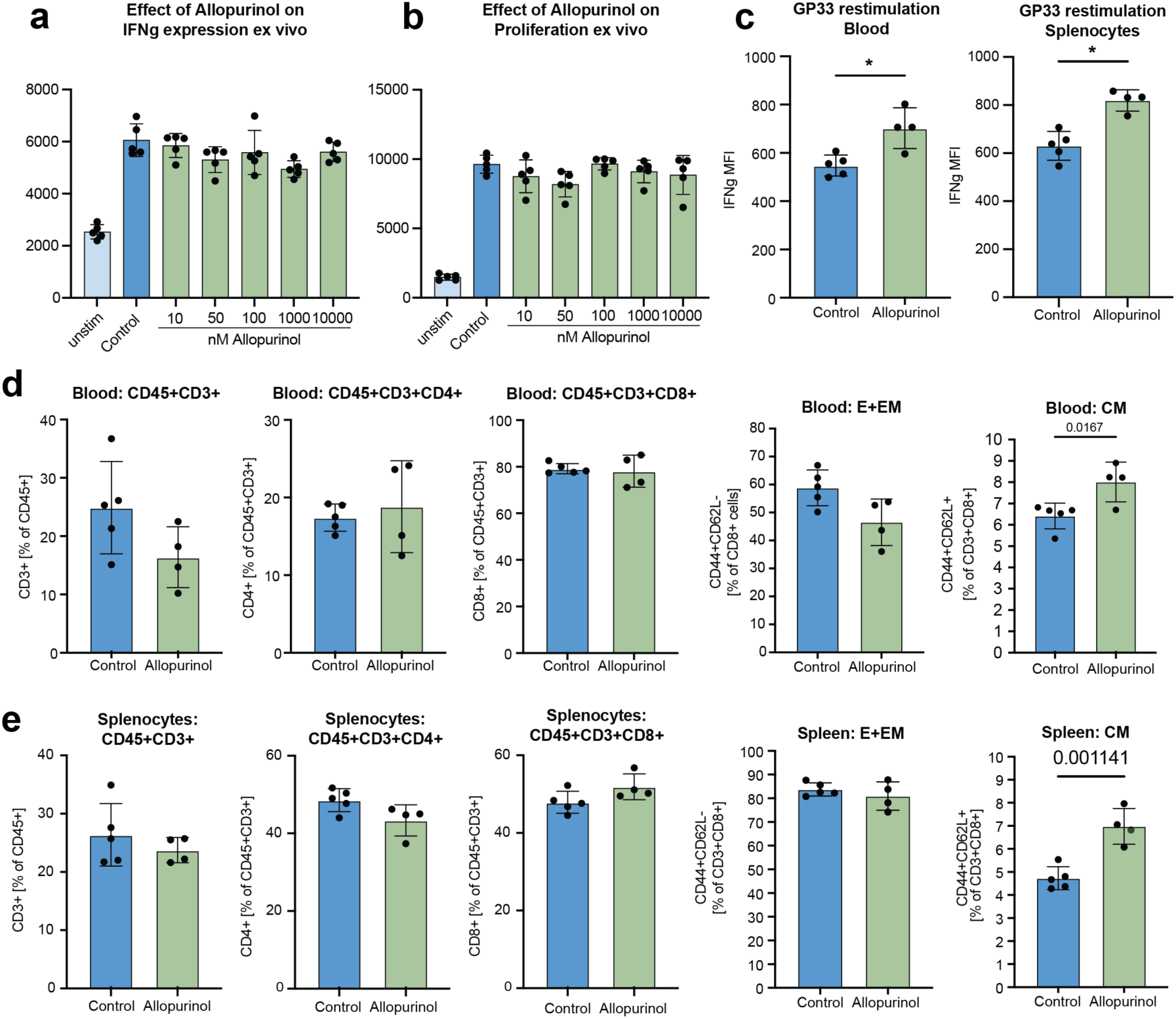
Elevation of serum purine nucleobase concentrations improves antiviral immune responses. a-b) High-throughput imaging-based analysis of the effect of different concentrations of allopurinol on **a)** effector function and **b)** proliferation of CD8^+^ T cells *in vitro*. Sample size is n = 5. **c)** Flow cytometric analysis of peptide restimulated CD8^+^ T cells at 8 days post LCMV Cl13 infection. Statistical analysis: Independent t-test. The graph represents a composite of two independent experimental replicates. **d-e)** Flow cytometry-based analysis of T cell subsets in **d)** blood and **e)** splenocytes. Statistical analysis: Independent t-test. The graph is a representative example of two independent experiments. Data in all subfigures are presented as mean ± SD with sample size n=5 in naïve and control or n=4 in treatment group, respectively.

**Extended Data Table 1: Viral infection metabolomics analysis.**

**Extended Data Table 2: Results of targeted metabolite screen.**

**Extended Data Table 3: Metabolomics analysis of metabolic treatments of CD8^+^ T cells.**

